# Simultaneous multimodal three-photon and optical coherence microscopy of the mouse brain in the 1700 nm optical window *in vivo*

**DOI:** 10.1101/2023.09.11.557176

**Authors:** Xusan Yang, Siyang Liu, Fei Xia, Meiqi Wu, Steven Adie, Chris Xu

## Abstract

Multimodal microscopy combining various imaging approaches can provide complementary information about tissue in a single imaging session. Here, we demonstrate a multimodal approach combining three-photon microscopy (3PM) and spectral-domain optical coherence microscopy (SD-OCM). We show that an optical parametric chirped-pulse amplification (OPCPA) laser source, which is the standard source for three-photon fluorescence excitation and third harmonic generation (THG), can be used for simultaneous OCM, 3-photon (3P) fluorescence and THG imaging. We validated the system performance in deep mouse brains *in vivo* with an OPCPA source operating at 1620 nm center wavelength. We visualized small structures such as myelinated axons, neurons, and large fiber tracts in white matter with high spatial resolution non-invasively using linear and nonlinear contrast at >1 mm depth in intact adult mouse brain. Our results showed that simultaneous OCM and 3PM at the long wavelength window can be conveniently combined for deep tissue imaging *in vivo*.

## Introduction

Optical microscopy is an essential tool for the non-invasive monitoring of biological structures and functions *in vivo*. Due to light scattering and absorption by biological tissue, it becomes challenging to obtain images with high contrast and high resolution in deep tissue, and the imaging depth in the mammalian brain is ultimately limited by the scattering of light despite the progress in the last three decades^1–3^. Because of the three-dimensionally confined excitation volume and the long excitation wavelength, two-photon (2P) microscopy achieved deep brain imaging at a depth of > 500 μm in practice in the adult mouse brain. Using 1280 nm excitation, a penetration depth of up to 1.6 mm can be achieved *in vivo* for imaging vasculature in the mouse neocortex with 2P excitation^4^. By carefully investigating the combined effects of the scattering and absorption of the brain tissue, the spectral windows of 1.3 *μ*m and 1.7 *μ*m^5,6^ have been shown to be optimum for deep brain imaging^7^, and long wavelength 3-photon microscopy (3PM) of neuronal structure and activity in the subcortical areas at >1-mm depth *in vivo* were demonstrated within these two wavelength windows. With specific fluorescence labels in different cell types, the structure and function of the cells, as well as cellular and sub-cellular dynamics, can be imaged quantitatively under a multiphoton fluorescence microscopy. Parallel to the development of fluorescence microscopy, a variety of approaches based on label-free contrast mechanisms, including reflectance confocal^8^, spectral reflectance confocal^9,10^, optical coherence microscopy^11– 14^, multi-harmonic generation^15,16^, Raman scattering^17,18^, photoacoustic microscopy^19^ have been developed in the last two to three decades. These modalities complement the exogenous fluorescence contrast mechanism.

Optical coherence tomography (OCT) combines coherence gating and spatial confocal gating, providing fast, non-invasive, and label-free imaging capability in tissues with penetration depths up to several millimeters. Compared to OCT, optical coherence microscopy (OCM) utilizes a higher numerical aperture (NA) objective and can provide cellular-resolution and label-free imaging of structures in the deep brain^12,20,21^. OCM can achieve a larger imaging depth than confocal microscopy by better rejecting multiply scattered and out-of-focus light^22^. By using a laser source in the 1.7 *μ*m window, researchers have achieved an imaging depth >1.2 mm in mouse brains^11,12,14^ with OCT and OCM.

Several groups reported combined scanning optical coherence and two-photon-excited fluorescence microscopy^23–26^ of skin and eye. However, the depth penetration capability of conventional 2P microscopy (2PM) is limited ^27,28^ and is typically well short of the imaging depth achievable by long-wavelength OCM. Since 3PM permits deeper imaging than 2PM and naturally requires long excitation wavelength, long-wavelength 3PM is a better choice than 2PM to combine with OCM for multimodal imaging in the deeper regions. An additional advantage of long wavelength 3PM is the accompanying label-free signal of THG. Indeed, THG and 3PM have been combined routinely in previous works^5,6,29^ for vasculature and neuronal imaging in the deep brain regions. Here we have developed a multimodal imaging system that combines OCM, THG, and 3PM to collect structural (OCM and THG) and molecularly specific (3P fluorescence) information simultaneously in a 3P laser scanning microscope. An OPCPA laser source, which is typically used for 3PM, was used for both 3PM and SD-OCM. We show high spatial resolution and high contrast imaging at > 1 mm in the intact adult mouse brain *in vivo* for both imaging modalities simultaneously. Our results demonstrate that multimodal imaging by combining SD-OCM and 3PM can provide complementary information deep within biological tissues.

## Results

One of the major criteria in developing the system for multimodal SD-OCM and 3P imaging (including 3P fluorescence and THG) is to preserve the performance of both imaging modalities, and to enable co-registered multimodal imaging with minimum additional complexity. The setup developed here is based on a typical multiphoton microscope where we combine SD-OCM and 3PM by sharing the same laser, scanning unit, and imaging optics.

### Multimodal microscopy setup combining OCM, THG and 3P imaging

Fig. 1(a) illustrates the multimodal microscopy setup based on a conventional laser scanning microscope design, where SD-OCM, THG, and 3P signals are collected in the epi-direction simultaneously. The major optical components of the multimodal microscopy setup are shown as two modules, the SD-OCM module and the 3P/THG module. The common source for the multimodal system is a wavelength-tunable optical parametric amplifier (OPA, Opera-F, Coherent) pumped by a femtosecond laser (Monaco, Coherent). A silicon wafer was employed to compensate for the dispersion of the optics in the laser source and the microscope, including the objective^30^ at the long excitation wavelength in the 1700 nm optical window. The laser repetition rate was maintained at 1 MHz for imaging. We set the center wavelength at 1620 nm, and the spectrum of the laser output is shown in the inset of Figure 1 (measured by a spectrometer, OSA207C, Thorlabs). The images were taken with a high-NA objective (Olympus XLPLN25XWMP2, 25×, NA 1.05), as shown in Fig. 1(a). The diameter of the illumination beam was measured to be 18.4 mm (1/e^2^ full width), which overfilled the back-aperture of the objective (∼15 mm)^31^. For the SD-OCM setup, the backscattered light was de-scanned, refocused by an aspherical lens (A110TM-C, Thorlabs) into the single-mode fiber (SMF-28, Thorlabs), and detected by a line CCD-based spectrometer (Wasatch Photonics, Cobra). All the images were simultaneously acquired by the data acquisition cards (NI PCI 6251, and the camera link card NI PCI-1424, National Instruments). The sensitivity of the OCM system was measured to be 96 dB using the characterization methods described in a previous publication^32^, with 2.1 mW average power and 1 MHz laser repetition rate. For the 3PM setup, two photomultiplier tubes (PMTs) with GaAsP photocathodes (H7422-40, Hamamatsu Photonics) were employed for the detection of the 3P fluorescence and THG signal. Transimpedance amplifiers (C7319, Hamamatsu) were used to convert the PMT current to voltage with the gain 10^6^ V/A for the fluorescence channel and 10^5^ V/A for the THG channel. All images from THG, 3PM and OCM were acquired by a custom LabVIEW software (National Instrument).

**Figure 1.**
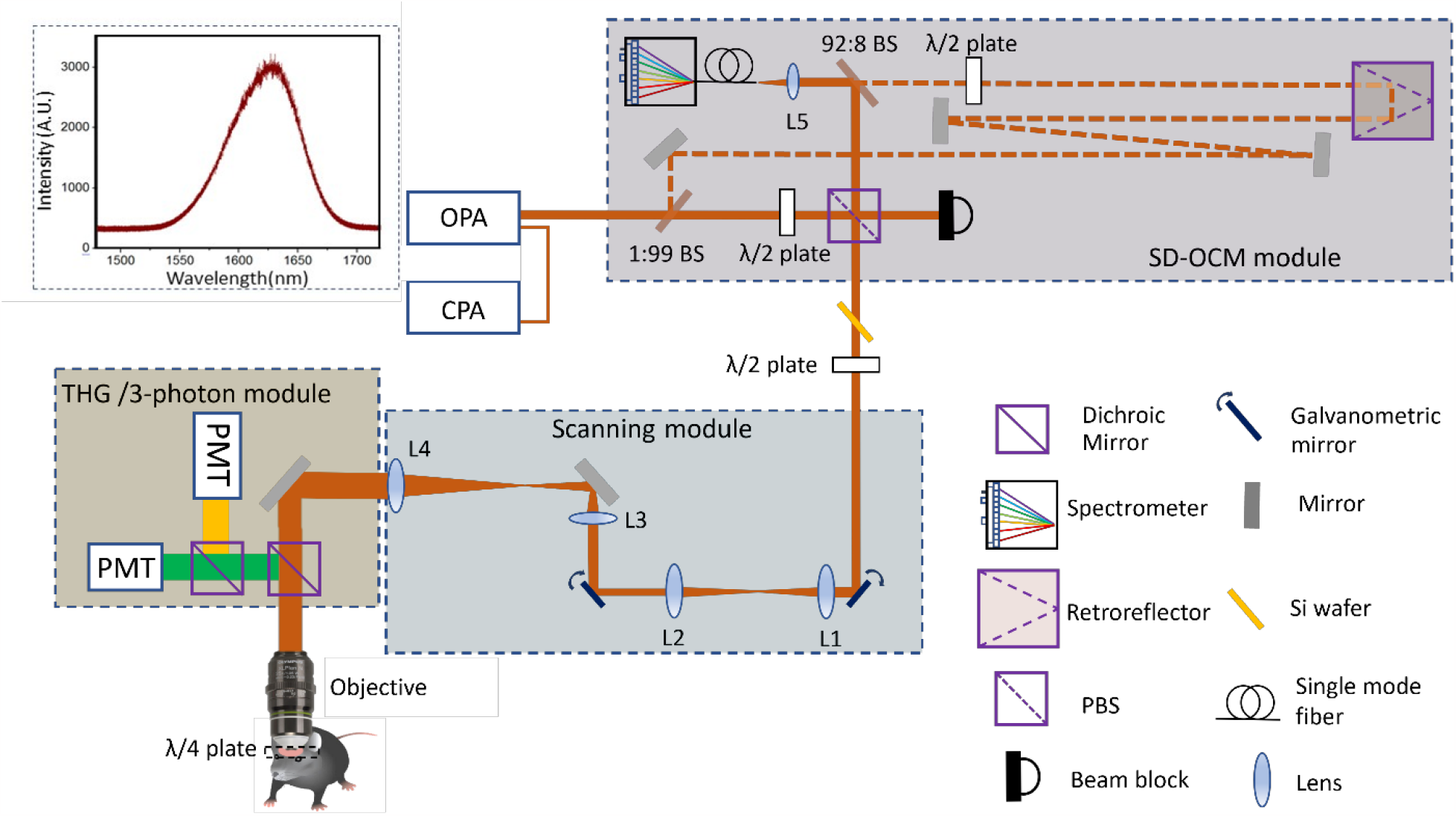
Setup of the multimodal microscopy system. The multimodal imaging system includes two paths, the OCM path and the THG/three-photon fluorescence path. The three modalities, OCM/ THG/3-photon, share the same laser source, the scanning unit and the imaging optics. The THG/3-photon path employed two PMTs, two dichroic mirrors, and two bandpass filters to separate the signals. In the OCM path, the reference arm and single-mode fiber-connected spectrometer were set up before the Si wafer for dispersion compensation. The scanning unit and signal collection system (two PMTs for THG and 3-photon, InGaAs line CCD-based high-speed spectrometer for OCM) are synchronized for signal collection under computer control. OPA: optical parametric amplifier. CPA: chirped-pulse amplifier. BS: beam splitter. PBS: polarizing beam splitter. PMT: photomultiplier tubes. Lens focal length: 150 mm for L1-L3, 750 mm for L4, and 6.24 mm for L5.

For deep brain imaging, we designed the multimodal microscopy setup to maximize the delivery of the illumination photons to the sample and the throughput of the epi-collected OCM signal photons. Although higher excitation power is needed for imaging deeper with 3PM, the reference arm only requires a fixed number of photons from the laser to generate an adequate spectrometer signal. Therefore, a 1:99 beam splitter (a thick glass slide) was employed to split the illumination power to the sample arm and the reference arm so that nearly all of the laser power was available for 3PM. Another 92:8 beam splitter (BP108, Thorlabs) preserves the signal strength of the backscattered photons for SD-OCM when re-combining the reference arm and the signal arm^33^. A λ/2 wave-plate and a PBS controlled the sample arm power, and a combination of λ/4 wave-plate as the mouse cranial window and λ/2 wave-plate in the sample arm ensures to optimize for high throughput for both the illumination and the backscattered photons at the PBS^8^ (see Methods for details).

### Characterization of the spatial resolutions deep in the mouse brain *in vivo*

In order to get simultaneous, co-registered information from each image voxel, closely matched spatial resolution in each modality is preferred, especially for the high-resolution OCM, THG, and 3-photon imaging shown in this work. Here we characterized our system resolution performance by measuring the small features in the mouse brain *in vivo*.

In our system, the OPCPA is tuned to a center wavelength of 1620 nm with a full-width-half maximum (FWHM) bandwidth of ∼70 nm, resulting in a theoretical coherence length of ∼11 μm. Since this is much larger than the confocal axial resolution (FWHM) (∼ 1.90 μm) associated with our high-NA (NA=1.05) objective, the axial resolution of our SD-OCM is determined by the objective lens and confocal detection. The theoretical FWHM axial resolution of our 3P imaging system is about 1.6 μm. The close match of the voxel sizes of the imaging modalities facilitates the calibration procedure to co-register the 3PM and OCM images in the axial direction and ensures the information in the images of the three imaging modalities were collected from approximately the same sample volume.

For all the imaging modalities, we used the smallest features (mainly the small capillaries and the axons) we could identify in each depth to estimate the resolution of the system (Fig. 2). Figs. 2h-k show that a high spatial resolution was achieved with all the three modalities with lateral resolutions ∼1 μm and axial resolutions ∼2.7 μm at the depth of ∼780 μm. We also measured the lateral FWHM of the small features at various depths and observed a gradual degradation of the lateral resolution as the imaging depth increases (Fig. 2i).

**Figure 2.**
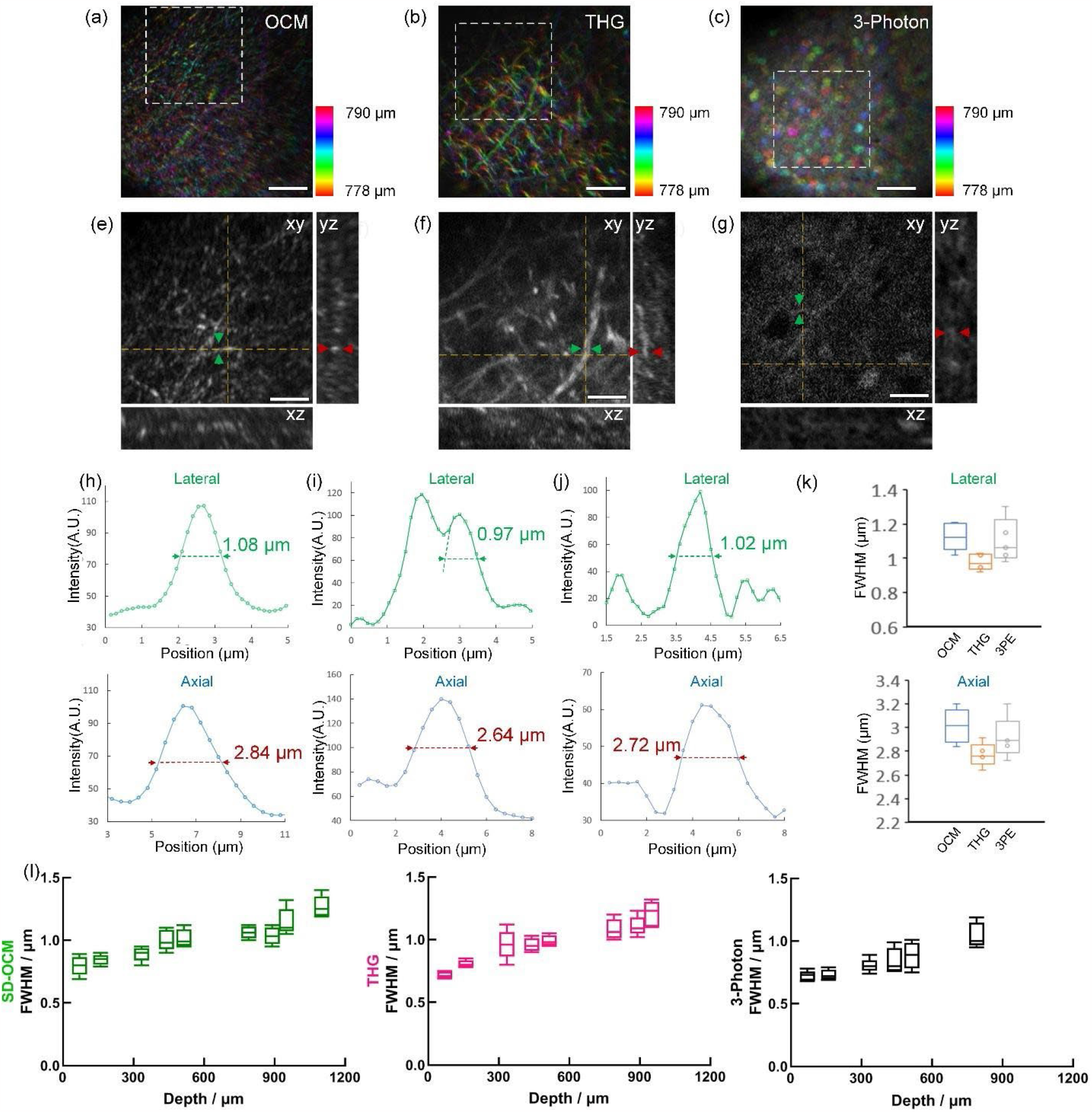
Resolution characterization of the multimodal imaging system in deep adult mouse brain *in vivo*. (a-c) OCM, THG, and 3P fluorescence imaging of mouse brain between 778 μm and 790 μm. The depth is coded by color. (e-g) Enlarged images of the boxes in (a-c) in three different projections. (h-i) The line profiles along the green and red arrows in (e-g.). (k) Boxplot of the lateral and axial resolution (N=5). (l) Boxplot of the lateral resolutions of SD-OCM, THG and 3-photon imaging measured at various imaging depths. Scale bars =50 μm.

### Image processing and co-registration

Co-registration of images from different modalities is required for multimodal imaging. In our system, OCM, THG, and 3PM shared the same laser source, scanning unit, and imaging optics in the illumination path, and the multimodal images were obtained simultaneously. Therefore, co-registration of the lateral positions of all three modalities was ensured. For the axial co-registration, the THG focus was first determined by maximizing the signal at the interface of the cover glass and the immersion water. The confocal pinhole position was then adjusted to maximize the OCM signal. This process ensures that the THG focal plane is conjugated to the OCM collection pinhole. In Figure Supplementary 1, the cross-section of the cover glass is shown to confirm the co-registration of OCM and THG.

Typical OCM uses telecentric scanning (Figure Supplementary 2a) in order to achieve uniform magnification across the field-of-view (FOV) at the cost of underfilling the objective lens. In our multimodal system, since only the image of the focal plane is taken, a non-telecentric scanning scheme (Figure Supplementary 2b) was used to overfill the objective lens. This maximizes illumination intensity for 3P fluorescence and THG. However, the non-telecentric scanning, which is routinely used in THG and 3P imaging, results in a field curvature in OCM because the focal plane of the objective does not correspond to the plane defined by the same optical path length^34^. Additionally, spherical aberration in the optical system of the sample arm (in particular the telescope lenses used for beam expansion) can lead to a variation of focal depth with transverse scan position. These effects can be compensated by estimating the focal plane curvature and removing it in post-processing, as shown in Figure Supplementary 2.

### Simultaneous *in vivo* multimodal imaging of mouse brain

*In vivo* simultaneous multimodal imaging of the mouse brain is performed to demonstrate the deep tissue imaging capability. Figure 4a shows the optical sections in mouse brain *in vivo* at various depths. The excitation powers at different depths were adjusted to achieve similar signal strength for all depths less than 1 mm. At depths beyond 1 mm, the maximum average power of 55 mW was used. The frame time was kept to a constant 3.9 seconds (515×512 pixels) for all the images taken.

**Figure 3.**
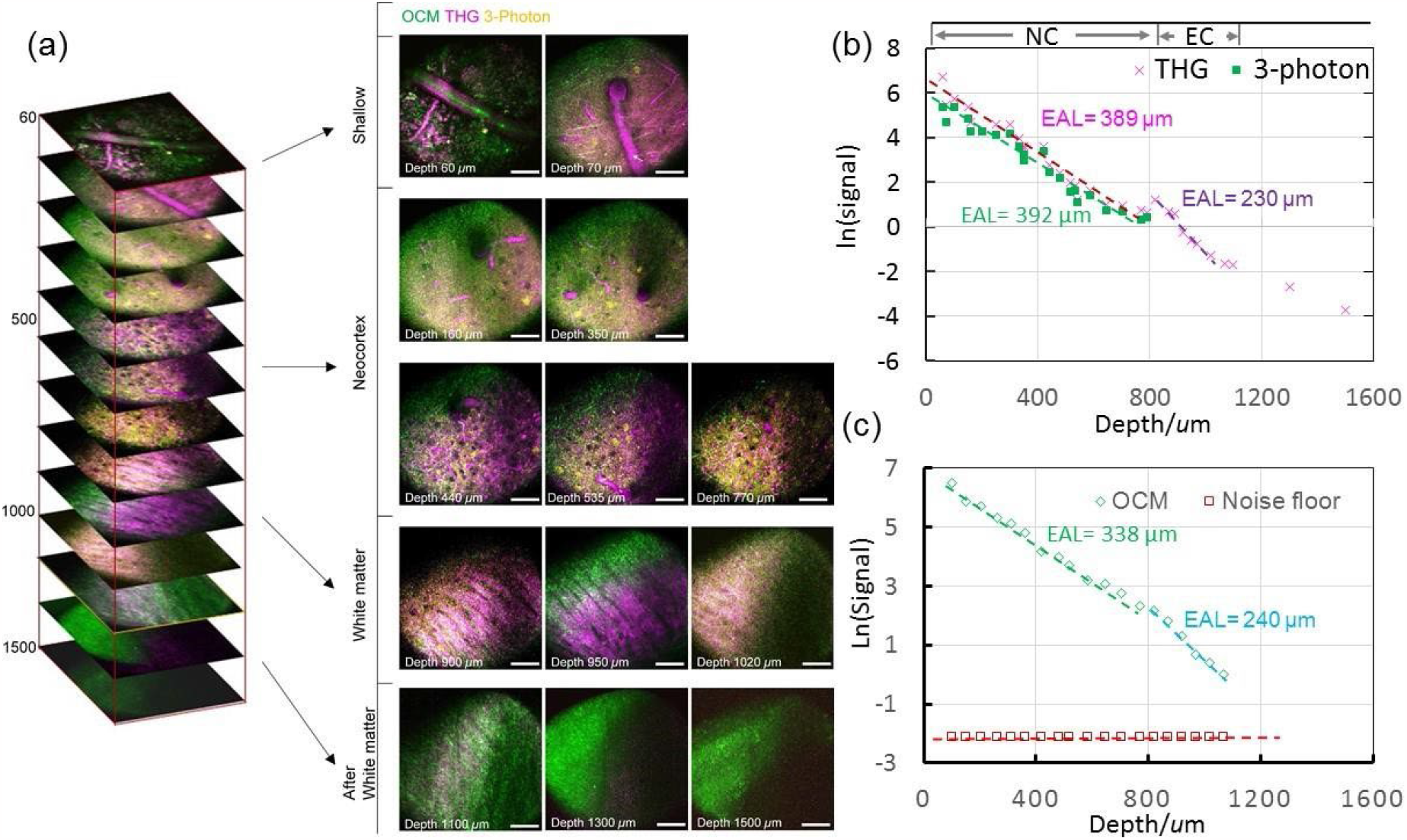
(a) Simultaneous multimodal imaging of the mouse brain *in vivo*. (b) THG and 3-photon signal attenuation curves of the neocortex (N.C.) and external capsule (E.C.) of mouse brain *in vivo*. The signal is normalized by the cube of the illumination power. (c) The signal attenuation curve and noise floor in SD-OCM are shown in green and red, respectively. The signal taken from the features of blood vessels is also normalized (Mag^2^/Power). Noise is taken away from the focus, and is also normalized (Mag^2^/the power at the surface). All frames before 1100 μm were individually normalized. Frames deeper than 1100 μm were normalized to the frame at 1100 μm. Scale bars = 50 μm.

**Figure 4.**
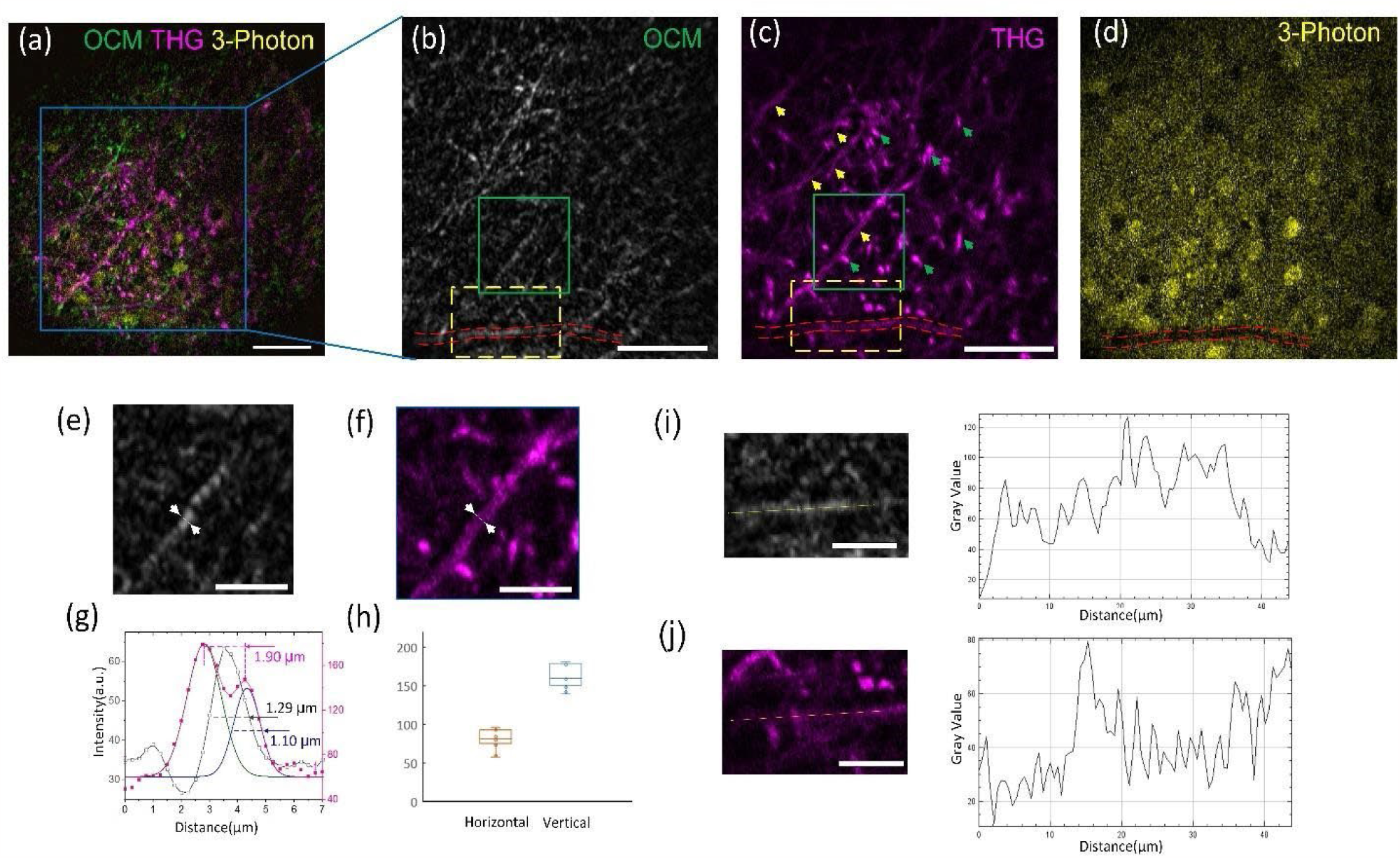
Multi-contrast multimodal imaging of neurons and myelinated axons. (a) Multimodal imaging at a depth of 784 *μ*m. (b-d) OCM, THG, and 3PM images of the boxed region in (a). (e) and (f) Enlarged views of the green boxes in (b) and (c), respectively. (g) Line profiles in dashed black and magenta corresponding to the lines in (e) and (f), respectively. The distance between the two peaks in the THG signal line profile (in magenta) is 1.9 *μ*m, which corresponds to the diameter of the myelin. The FWHM of the Gaussian fit for the right peak of the THG signal line profile is 1.1 *μ*m, highlighting the capability of THG to directly identify fine features in myelin. The FWHM of the OCM signal line profile (dashed black line) is 1.29 *μ*m. (h) THG signal intensity (a.u.) of horizontal and vertical myelin in (c). (i) and (j). Left, enlarged views of the yellow boxes in (b) and (c), respectively; right, line profile corresponding to dashed lines in the left panels. Scale bars in (a) and (c): 50 *μ*m. Scale bars in (e, f, i, j), 20 *μ*m.

The three-photon excited fluorescence and THG signal within the focal volume is mostly generated by ballistic photons, while the collected signal includes both ballistic photons and multiply scattered photons in THG and 3PM. From Figure 3, it is possible to estimate the effective attenuation length (EAL), a parameter combining the scattering and absorption length of the tissue, for 1620 nm imaging in mouse brain tissue *in vivo*. For THG and 3PM, we take the average values of the blood vessels and neuron soma, respectively, as the THG and 3P signals, and we estimate the EAL as the depth where the signal attenuates by 1/e^3^. Figure 4b shows the detected THG signal (in purple) as a function of imaging depth. The EAL is determined to be 389 μm between 60 μm and 800 μm depth in the neocortex of the mouse brain. The EAL is ∼230 μm between 800 μm and 1000 μm depth in the external capsule. From the data of 3P fluorescence signal versus depth (in green, mostly from neuron somas), the EAL between 60 μm and 800 μm depth is approximately 392 μm.

Both the signal generation and collection within the focal volume in SD-OCM are mostly from ballistic photons. Therefore, ballistic photon attenuations in both the illumination and collection paths need to be taken into count in the SD-OCM case when measuring the tissue attenuation length in the mouse brain. For the SD-OCM data acquisition, the reference arm position was adjusted for each focus depth to position the focus at a fixed optical path delay, in order to avoid the sensitivity roll-off from the spectrometer. We measured the blood vessel OCM signal versus depth (Fig. 3c), and compared it with the *en face* frame-averaged signal decay in Supplementary Fig. 3. We found that the signal of a specific feature (e.g., the blood vessel) versus depth provides a better estimate of the EAL than the frame-averaged signal, which includes all features and is highly dependent on the brightest features at each depth. The OCM noise floor was estimated from the average intensity in the out-of-focus depths. Note that the OCM noise floor is flat because the power in the reference arm was kept at a constant level and was much higher than the signal power from the sample arm at all depths.

### Multimodal contrast for deep mouse brain imaging *in vivo*

Multimodal cellular imaging combining 3PM, THG, and SD-OCM can provide multiple unique and complementary contrasts in the mouse brain *in vivo*. Here we investigate the similarities and differences of these contrasts.

### Neurons and myelinated axons visualized by multimodal imaging

Fig. 4 shows 3P, THG and OCM images simultaneously obtained in the deep cortical layers at the depth of 784 *μ*m, and illustrates the complementary information with both labeled and label-free contrast, such as the neurons in the 3P fluorescence channel and myelinated axons in the THG and OCM channels. 3PM with fluorescence labeling strategies such as genetic targeting provides high specificity information which is not accessible with OCM and THG imaging. In Supplementary Figure 4, neurons were observed with bright signals by 3PM while they were not visible in the OCM and THG images. For OCM, the main source of image contrast comes from the linear refractive index variation. In the mouse brain, the myelinated axons have a high refractive index^9,35^ due to the high lipid content and exhibit a high scattering contrast. For THG signal, it is sensitive to the nonlinear refractive index variations and the geometry of the features^16^. In Fig. 4c, the interface of the myelinated axons and the surroundings can be visualized by THG imaging but not OCM. As shown in Figure 4g, the THG signal of the myelinated axons is the strongest from the sides of the myelin tube and exhibits two peaks, while the OCM signal is mainly generated from the top and bottom of the myelin tube and has a single peak. Using the THG signal, the diameter of the myelin can be measured as 1.9 *μ*m. Both horizontal and vertical interfaces in the focal volume will elicit strong THG signal, and the vertical interface (green arrows in Figure 4c) generates a higher THG signal than the horizontal one (yellow arrows in Figure 4c)^36,37^ (see Figure 4h), while the vertically oriented myelins are mostly not visible in the OCM images.

We found that OCM images of myelin show a patchy, fragmented structure. This can be attributed to self-interference between the top and bottom interfaces of myelin, similar to thin-film interference^38^. This distinctive pattern can help with identifying myelin in the deep mouse brain *in vivo*. For small blood vessel, the dark streak or bright line patterns of the flowing red blood cells (RBCs) in vessels were revealed by 1700 nm^39^ and 1300 nm^40^ THG imaging. Here stripe patterns show up in both THG and OCM signal (RBCs, the dotted lines in Figure 4i and j). The blood vessels are not fluorescently labeled and appear as dark shadows (Figure 4d) in 3P fluorescence images.

Figure 5a shows the three-dimensional reconstruction of large horizontal blood vessels in the shallow regions of the mouse brain by OCM and THG imaging. In both SD-OCM and THG images, we can visualize the intensity fluctuations across the blood vessel caused by the motion blur of moving RBCs (Figure 5b, 5c). In the OCM image, the middle of the blood vessel generated a stronger signal than the two sides, whereas THG signal levels were similar for both center and sides. Therefore, OCM signals have a smaller FWHM than THG (Fig. 5e). We also observed differences in signal from blood vessels of different sizes (see Figure Supplementary 5 and 6). Vertical blood vessels appear dark compared to the surroundings in OCM (Figures 5f and Supplementary 7a); however, the corresponding THG signal is stronger due to the contributions by the RBCs. The blood vessel walls, both vertical and horizontal, generate strong THG signals, while they produce weak or no detectable OCM signals (Figures 5b and 5f). The vessel wall thickness, as estimated from the THG image (the right panels in Figure 5f and Supplementary Figure 7), is ∼2 *μ*m. This is consistent with the vessel wall thickness of arterioles or venules in the mouse brain^43^.

**Figure 5.**
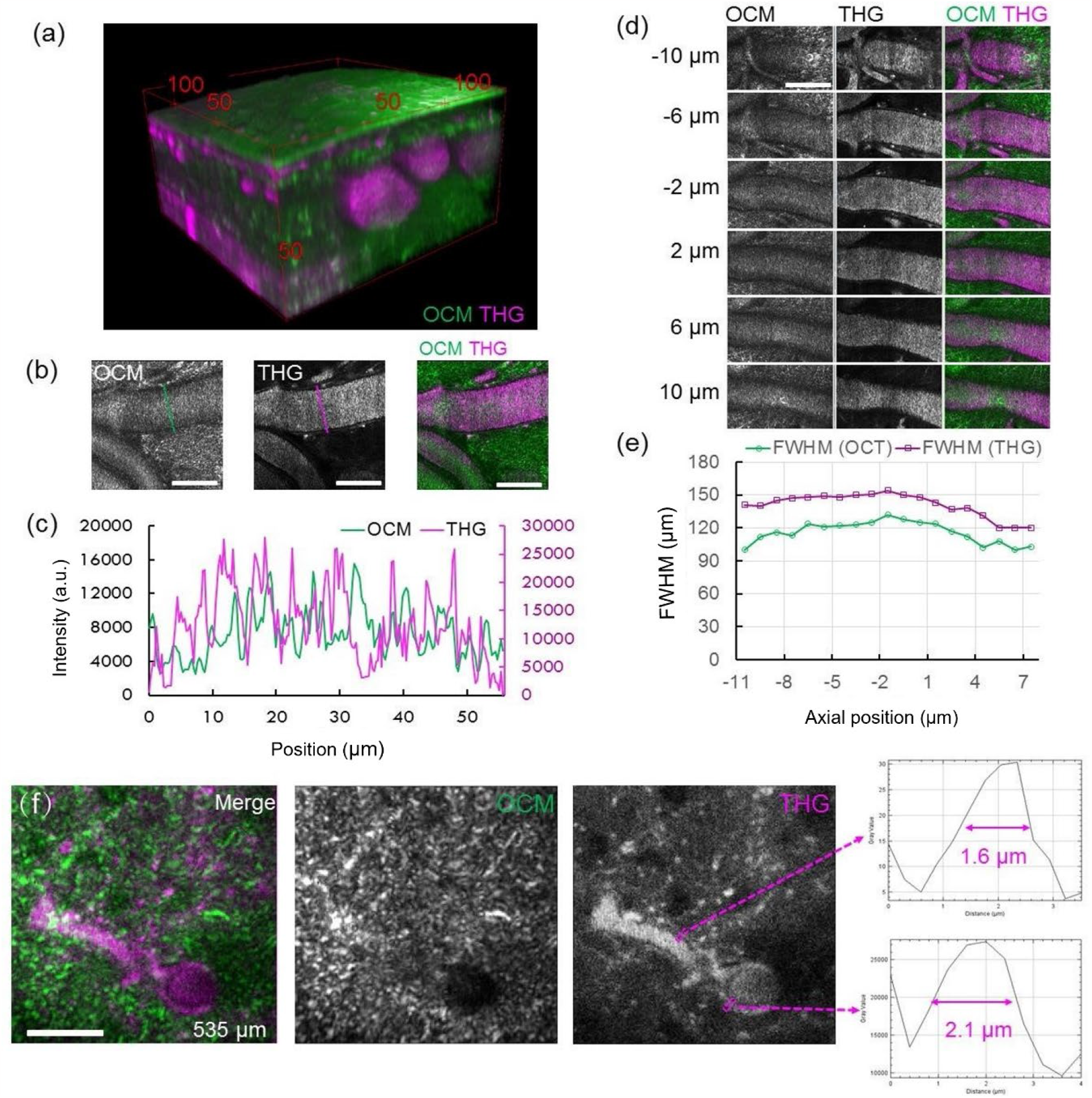
Multi-contrast of large blood vessels in mouse brain by multimodal imaging. (a) Three-dimensional reconstruction of multimodal images from a mouse brain at depth up to 80 *μ*m. All frames were individually normalized. (b) Normalized frames of OCM, THG, and merged signals at a depth of 34 *μ*m. OCM and THG profiles of the green and magenta dotted lines across the vessels are displayed in (c). (d) Normalized frames of the OCM, THG, and merged signals of large blood vessels at various depths (the center of the blood vessel was set at 0 *μ*m and the depths of the other images are measured relative to the center). (e) Plot of OCM and THG signals FWHM values corresponding to (d). (f) Normalized frames of OCM, THG, and merged signals of a vertical blood vessel and a horizontal one at a depth of 535 *μ*m. The line intensity profiles crossing the blood vessel walls in the THG image are plotted on the right. Scale bars in (b) and (c), 50 *μ*m; scale bar in (f), 20 *μ*m.

## Conclusions and prospective

In this paper, we demonstrated the use of a single OPCPA laser source, commonly used in three-photon microscopy (3PM), for simultaneous OCM, THG, and three-photon excited fluorescence imaging, and performed multimodal, high spatial resolution imaging of the intact adult mouse brain *in vivo* at depth > 1 mm. 3PM is emerging as a valuable tool for studying neuronal structure and function using exogenous dyes or fluorescent proteins because of its capability for deep tissue penetration in the mouse brain. Meanwhile, OCM and THG could provide unique and complementary label-free contrast of brain features such as blood vessels, blood cells, and myelinated axons. Therefore, the multiple contrast mechanisms of these three modalities could offer complementary structural and physiological information.

The phase information from OCM can be potentially utilized in the future for aberration sensing ^41,42^. The wavefront information provided by OCM can then be used to perform hardware adaptive optics for aberration correction. Such closed-loop aberration sensing and correction can be especially beneficial for 3PM because the higher-order nonlinear excitation is more sensitive to wavefront imperfections^42^. Therefore, the combination of 3PM and SD-OCM offers not only the linear and the nonlinear contrast from the sample simultaneously, but also the intriguing possibility of using the information obtained in one modality to enhance the imaging performance of the other.

### Online Methods Mouse preparation

All animal procedures were reviewed and approved by the Cornell Institutional Animal Care and Use Committee. We used male B6.Cg-Tg(Thy1-Brainbow1.0)HLich/J mice (eight weeks old, The Jackson Laboratory) for red fluorescence protein (RFP)-labelled neuron imaging. Animals were prepared using the methods described in the previous work^8, 44^. We used a 5 × 5 mm square-shaped λ/4 waveplate (WPQ501, Thorlabs) as the cranial window instead of the normal cover glass. Craniotomies were centered at 2.5 mm posterior and 2 mm lateral to the Bregma point. As the mice were not subjected to Cre recombinase, RFP tdimer2(12) was the only fluorescent protein expressed.

### *In vivo* imaging of mouse brain

We imaged RFP-labeled neurons in a B6.Cg-Tg(Thy1-Brainbow1.0)HLich/J mouse two weeks after surgery. The mouse was mounted on a motorized stage (M-285, Sutter Instrument Company) for axial scanning. Deuterium oxide (i.e. heavy water, D_2_O) was used as the objective immersion medium to reduce the absorption caused by water (H_2_O) when imaging at 1620 nm. Because the index of refraction of brain tissue is larger than that of the immersion medium (heavy water), the actual imaging depth within the tissue was ∼5–10% more than indicated by the depth measurement (i.e. the raw axial movement of the objective).

### OCM image processing

SD-OCM images were reconstructed from spectral data through the following steps: wavelength to wavenumber domain resampling, reference background subtraction, dispersion compensation between sample and reference arm, and finally a Fourier transform along wavenumber to convert data from spectral to space domain. In addition, sample tilt and coherence gate curvature were compensated by adding a linear phase ramp in the spectral domain for each A-scan (Fig.S2). The slope of the phase ramp at each transverse location was estimated by fitting a second-order polynomial to the cover glass surface. When averaging, a location was exposed for an extended amount of time, with the chance that *in vivo* biological motion such as heartbeats may corrupt the averaged results. In such cases, a bulk shift based on cross-correlation to a reference image was applied to eliminate the motion across time.

## Acknowledgements

This research is funded by National Science Foundation (Career: CBET-1752405, DBI-1707312); National Institutes of Health (NIBIB-R21EB022927, NINDS-R01NS120819). The authors acknowledge Dr. Yusaku Hontani for proofreading the manuscript.

## References

1. Denk, W., Strickler, J. H. & Webb, W. W. Two-photon laser scanning fluorescence microscopy. Science (80-.). (1990) doi:10.1126/science.2321027.

2. Lichtman, J. W. & Denk, W. The big and the small: Challenges of imaging the brain’s circuits. Science (2011) doi:10.1126/science.1209168.

3. Helmchen, F. & Denk, W. Deep tissue two-photon microscopy. Nature Methods (2005) doi:10.1038/nmeth818.

4. Kobat, D. et al. Deep tissue multiphoton microscopy using longer wavelength excitation. Opt. Express (2009) doi:10.1364/oe.17.013354.

5. Horton, N. G. et al. In vivo three-photon microscopy of subcortical structures within an intact mouse brain. Nat. Photonics (2013) doi:10.1038/nphoton.2012.336.

6. Ouzounov, D. G. et al. In vivo three-photon imaging of activity of GcamP6-labeled neurons deep in intact mouse brain. Nat. Methods (2017) doi:10.1038/nmeth.4183.

7. Wang, M. et al. Comparing the effective attenuation lengths for long wavelength in vivo imaging of the mouse brain. Biomed. Opt. Express (2018) doi:10.1364/boe.9.003534.

8. Xia, F. et al. In vivo label-free confocal imaging of the deep mouse brain with longwavelength illumination. Biomed. Opt. Express (2018) doi:10.1364/boe.9.006545.

9. Schain, A. J., Hill, R. A. & Grutzendler, J. Label-free in vivo imaging of myelinated axons in health and disease with spectral confocal reflectance microscopy. Nat. Med. (2014) doi:10.1038/nm.3495.

10. Kwon, J. et al. Label-free nanoscale optical metrology on myelinated axons in vivo. Nat. Commun. (2017) doi:10.1038/s41467-017-01979-2.

11. Chong, S. P. et al. Noninvasive, in vivo imaging of subcortical mouse brain regions with 17 μm optical coherence tomography. Opt. Lett. 40, 4911 (2015).

12. Yamanaka, M., Teranishi, T., Kawagoe, H. & Nishizawa, N. Optical coherence microscopy in 1700 nm spectral band for high-resolution label-free deep-tissue imaging. Sci. Rep. (2016) doi:10.1038/srep31715.

13. Wang, H. et al. Polarization sensitive optical coherence microscopy for brain imaging. Opt. Lett. (2016) doi:10.1364/ol.41.002213.

14. Park, K. S., Shin, J. G., Qureshi, M. M., Chung, E. & Eom, T. J. Deep brain optical coherence tomography angiography in mice: in vivo, noninvasive imaging of hippocampal formation. Sci. Rep. 8, 1–7 (2018).

15. Lim, H. et al. Label-free imaging of Schwann cell myelination by third harmonic generation microscopy. Proc. Natl. Acad. Sci. U. S. A. (2014) doi:10.1073/pnas.1417820111.

16. Witte, S. et al. Label-free live brain imaging and targeted patching with third-harmonic generation microscopy. Proc. Natl. Acad. Sci. U. S. A. (2011) doi:10.1073/pnas.1018743108.

17. Wang, H., Fu, Y., Zickmund, P., Shi, R. & Cheng, J. X. Coherent anti-stokes Raman scattering imaging of axonal myelin in live spinal tissues. Biophys. J. (2005) doi:10.1529/biophysj.105.061911.

18. Freudiger, C. W. et al. Label-free biomedical imaging with high sensitivity by stimulated raman scattering microscopy. Science (80-.). (2008) doi:10.1126/science.1165758.

19. Yao, J. et al. High-speed label-free functional photoacoustic microscopy of mouse brain in action. Nat. Methods 12, 407–410 (2015).

20. Yamanaka, M., Hayakawa, N. & Nishizawa, N. Signal-to-background ratio and lateral resolution in deep tissue imaging by optical coherence microscopy in the 1700 nm spectral band. Sci. Rep. 9, 16041 (2019).

21. Srinivasan, V. J., Radhakrishnan, H., Jiang, J. Y., Barry, S. & Cable, A. E. Optical coherence microscopy for deep tissue imaging of the cerebral cortex with intrinsic contrast. Opt. Express (2012) doi:10.1364/oe.20.002220.

22. Izatt, J. A., Swanson, E. A., Fujimoto, J. G., Hee, M. R. & Owen, G. M. Optical coherence microscopy in scattering media. Opt. Lett. (1994) doi:10.1364/ol.19.000590.

23. Beaurepaire, E., Moreaux, L., Amblard, F. & Mertz, J. Combined scanning optical coherence and two-photon-excited fluorescence microscopy. Opt. Lett. (1999) doi:10.1364/ol.24.000969.

24. Tang, S., Krasieva, T. B., Chen, Z. & Tromberg, B. J. Combined multiphoton microscopy and optical coherence tomography using a 12-fs broadband source. J. Biomed. Opt. (2006) doi:10.1117/1.2193428.

25. Chong, S. P., Lai, T., Zhou, Y. & Tang, S. Tri-modal microscopy with multiphoton and optical coherence microscopy/tomography for multi-scale and multi-contrast imaging. Biomed. Opt. Express (2013) doi:10.1364/boe.4.001584.

26. Vinegoni, C. et al. Integrated structural and functional optical imaging combining spectral-domain optical coherence and multiphoton microscopy. Appl. Phys. Lett. (2006) doi:10.1063/1.2171477.

27. Takasaki, K., Abbasi-Asl, R. & Waters, J. Superficial bound of the depth limit of twophoton imaging in mouse brain. eNeuro (2020) doi:10.1523/ENEURO.0255-19.2019.

28. Theer, P. & Denk, W. On the fundamental imaging-depth limit in two-photon microscopy. J. Opt. Soc. Am. A (2006) doi:10.1364/josaa.23.003139.

29. Wang, T. et al. Three-photon imaging of mouse brain structure and function through the intact skull. Nat. Methods (2018) doi:10.1038/s41592-018-0115-y.

30. Horton, N. G. & Xu, C. Dispersion compensation in three-photon fluorescence microscopy at 1,700 nm. Biomed. Opt. Express 6, 1392–1397 (2015).

31. Bumstead, J. R. et al. Designing a large field-of-view two-photon microscope using optical invariant analysis. Neurophotonics 5, 25001 (2018).

32. Leitgeb, R., Hitzenberger, C. & Fercher, A. Performance of fourier domain vs time domain optical coherence tomography. Opt. Express (2003) doi:10.1364/oe.11.000889.

33. Auksorius, E. Light-efficient beamsplitter for Fourier-domain full-field optical coherence tomography. Opt. Lett. (2020) doi:10.1364/ol.383823.

34. Kumar, A. et al. Anisotropic aberration correction using region of interest based digital adaptive optics in Fourier domain OCT. Biomed. Opt. Express 6, 1124–1134 (2015).

35. Allegra Mascaro, A. L. et al. Label-free near-infrared reflectance microscopy as a complimentary tool for two-photon fluorescence brain imaging. Biomed. Opt. Express (2015) doi:10.1364/boe.6.004483.

36. Weigelin, B., Bakker, G. J. & Friedl, P. Third harmonic generation microscopy of cells and tissue organization. J. Cell Sci. (2016) doi:10.1242/jcs.152272.

37. Genthial, R. et al. Third harmonic generation imaging and analysis of the effect of low gravity on the lacuno-canalicular network of mouse bone. PLoS One 14, e0209079 (2019).

38. Binding, J. et al. Brain refractive index measured in vivo with high-NA defocus-corrected full-field OCT and consequences for two-photon microscopy. Opt. Express (2011) doi:10.1364/oe.19.004833.

39. Liu, H. et al. In vivo deep-brain blood flow speed measurement through third-harmonic generation imaging excited at the 1700-nm window. Biomed. Opt. Express 11, 2738–2744 (2020).

40. Ahn, S. J. et al. Label-free assessment of hemodynamics in individual cortical brain vessels using third harmonic generation microscopy. Biomed. Opt. Express 11, 2665–2678 (2020).

41. Adie, S. G., Graf, B. W., Ahmad, A., Carney, P. S. & Boppart, S. A. Computational adaptive optics for broadband optical interferometric tomography of biological tissue. Proc. Natl. Acad. Sci. U. S. A. (2012) doi:10.1073/pnas.1121193109.

42. Sinefeld, D., Paudel, H. P., Ouzounov, D. G., Bifano, T. G. & Xu, C. Adaptive optics in multiphoton microscopy: comparison of two, three and four photon fluorescence. Opt. Express 23, 31472–31483 (2015).

43. Müller, B., Lang, S., Dominietto, M., Rudin, M., Schulz, G., Deyhle, H., Germann, M., Pfeiffer, F., David, C. and Weitkamp, T.,. High-resolution tomographic imaging of microvessels. In Developments in X-ray tomography VI (Vol. 7078, pp. 89–98). SPIE (2008).

44. Xia, F., Gevers, M., Fognini, A., Mok, A.T., Li, B., Akbari, N., Zadeh, I.E., Qin-Dregely, J. and Xu, C. Short-wave infrared confocal fluorescence imaging of deep mouse brain with a superconducting nanowire single-photon detector. ACS Photonics, 8(9), pp.2800–2810 (2021).

