## Supplementary Information for "Simultaneous multimodal three-photon and optical coherence microscopy of the mouse brain in the 1700 nm optical window *in vivo*"

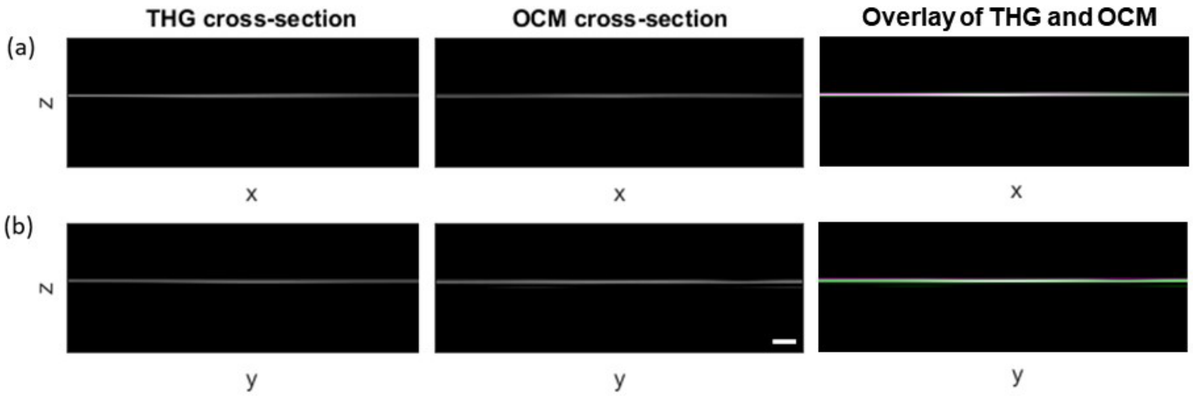


**Figure Supplementary 1.** Axial co-registration of THG and OCM signals. (a,b) XZ and YZ images of THG microscopy and OCM of a cover glass and the combined THG and OCM images (THG in magenta and OCM in green). THG and OCM peak signals appeared at the same depth in both XZ and YZ images. Scale bar represents 10 μm. All panels have the same field-of-view.

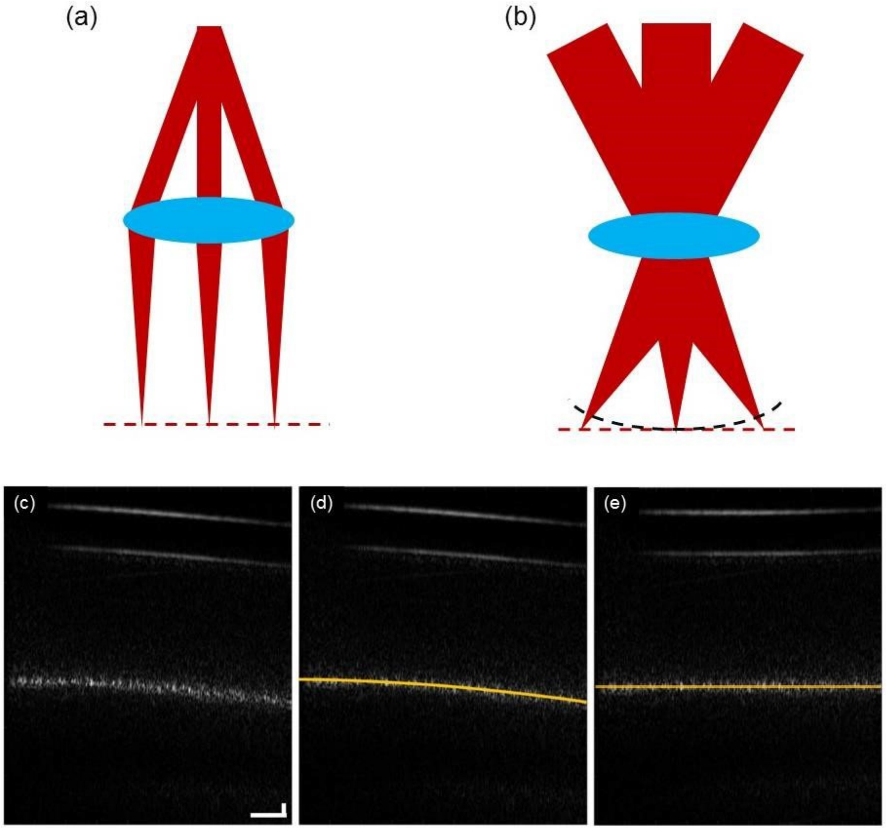


**Figure Supplementary 2. OCM curvature correction.** (a) Schematic for a telecentric scanning system in conventional OCT. (b) Schematic for a simplified 4-F conjugated non-telecentric scanning system in OCM. The dotted red line shows the flat focal plane, and the dashed black curve shows the curved coherence gate plane. (c) Raw data of an uncorrected OCM frame. (d) Fitting of a parabola to the focal plane curvature to estimate the required correction. (e) The corrected OCM frame.

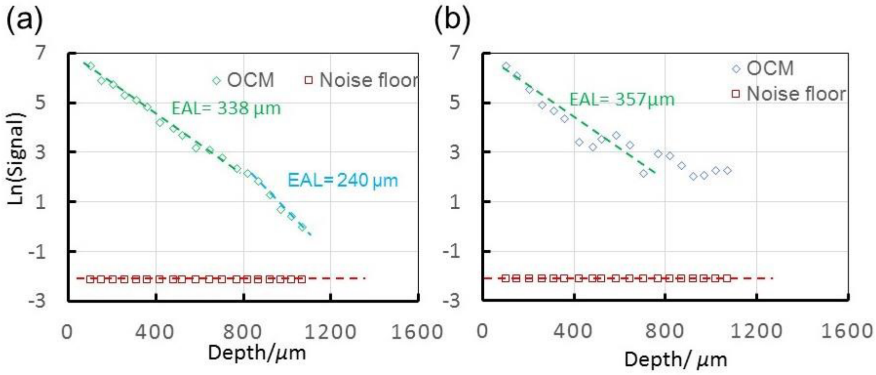


**Figure. Supplementary 3.** SD-OCM signal decay and noise floor. The signal attenuation curve and noise floor in SD-OCM are shown in green/light blue and red, respectively. The signal is taken from blood vessel features (a) and the frame average (b). Noise is taken at locations away from the focus, and is also normalized ($Mag^2$/the power at the surface). All frames before 1100 μm were individually normalized. Frames deeper than 1100 μm were normalized to the frame at 1100 μm.

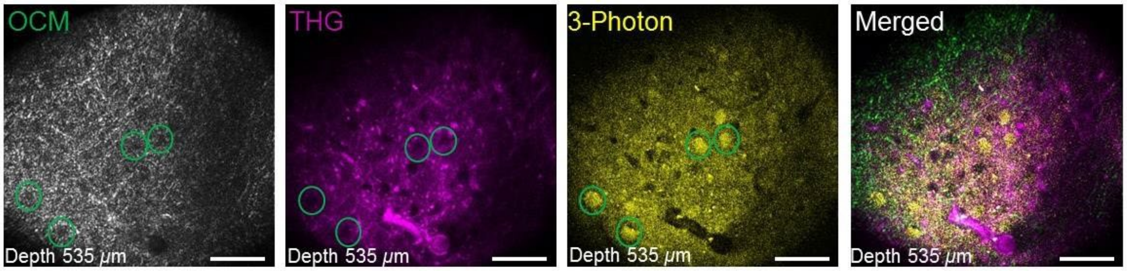


**Figure Supplementary 4. Multi-contrast of low-signal neuron "holes" by multimodal imaging**. Scale bar, 50 μm

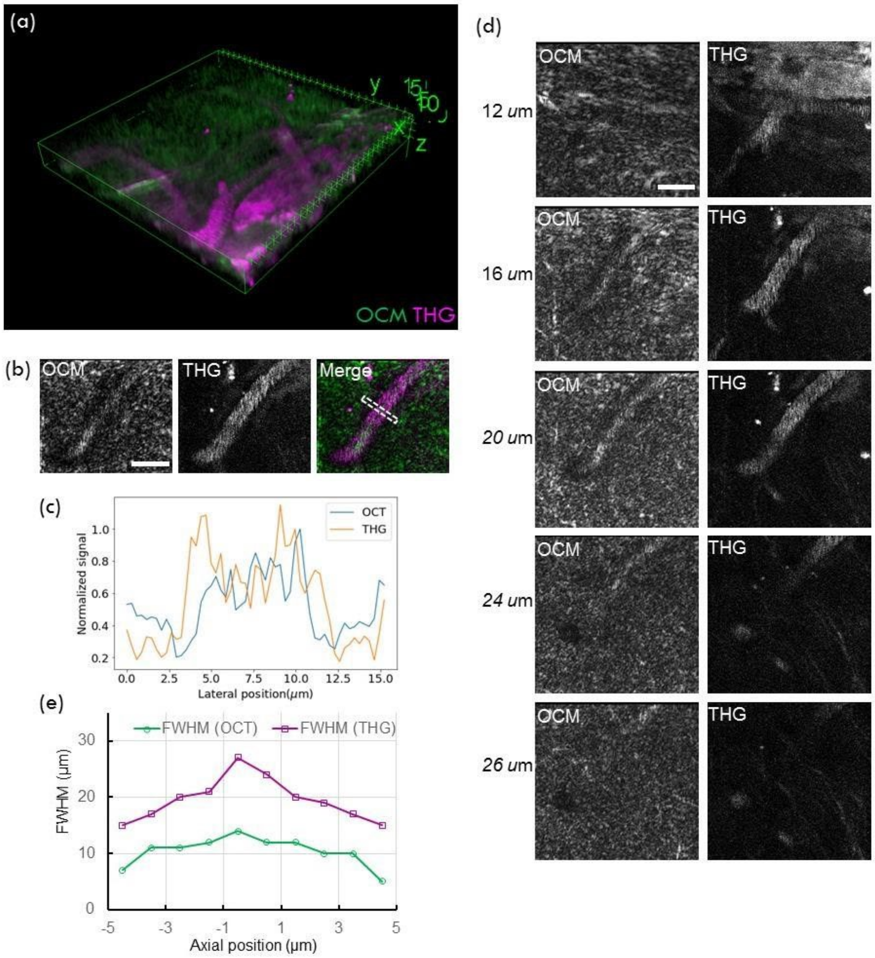


**Figure Supplementary 5. Multi-contrast of blood vessels in mouse brain by multimodal imaging**. (a) Three-dimensional reconstruction of multimodal images of a mouse brain. All frames were individually normalized. (b) Normalized frames of the OCM, THG, and merged signals at a depth of 19 µm. (c) OCM and THG profiles of the green and magenta dotted lines across the vessels of (b). (d) Normalized frames of the OCM, THG, and merged signals of large blood vessels at various depths. (e) FWHM values corresponding to (d). Scale bar in (b) and (c), 20 µm.

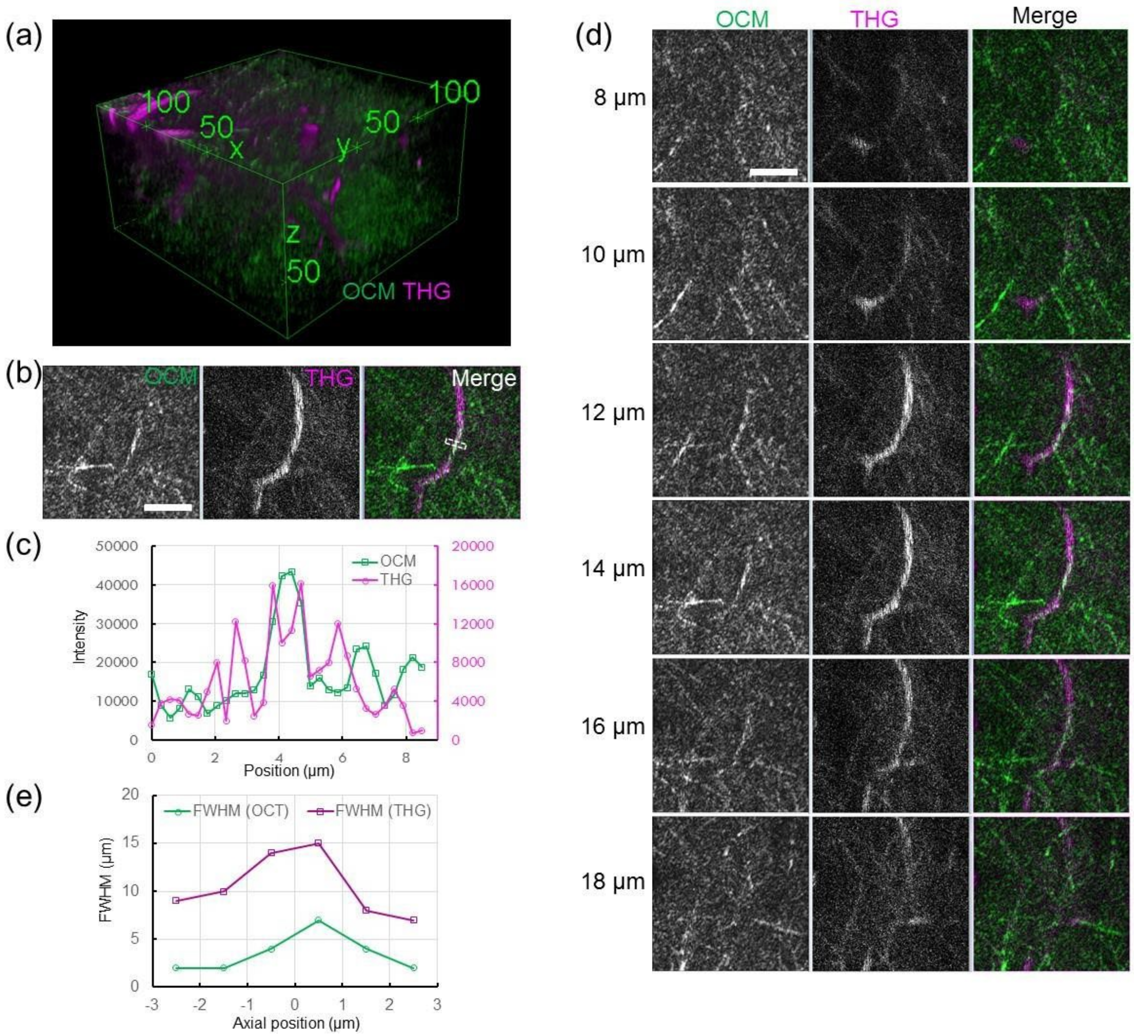


**Figure. Supplementary 6. Multi-contrast of medium-size blood vessels in mouse brain by multimodal imaging**. (a) Three-dimensional reconstruction of multimodal images of a mouse brain. All frames were individually normalized. The units for the numbers are in μm. (b) Normalized frames of the OCM, THG, and merged signals at a depth of 14 μm. (c) OCM and THG profiles of the green and magenta dotted lines across the vessels of (b). (d) Normalized frames of the OCM, THG, and merged signals of medium blood vessels at various depths. (e) FWHM values corresponding to (d). Scale bar in (b) and (c), 20 μm.

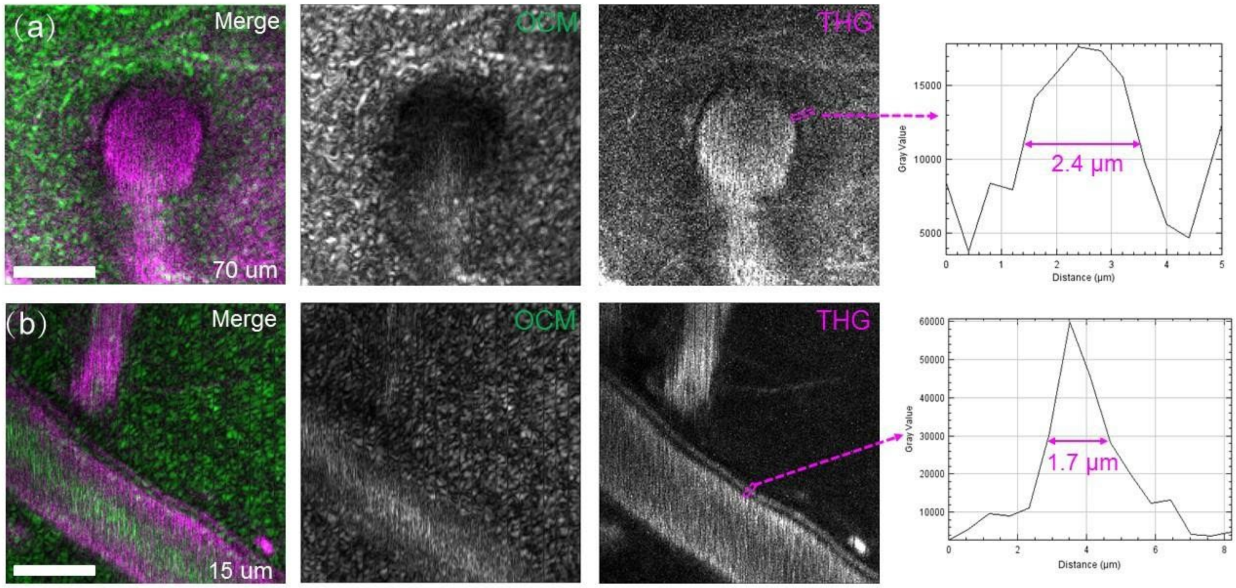


**Figure Supplementary 7. Multi-contrast of the blood vessel wall in mouse brain by multimodal imaging**. Normalized frames of the OCM, THG, and merged signals of blood vessels having different orientations with respect to the optical axis: (a) a large vertical blood vessel at a depth of 70 µm below the surface and (b) a large horizontal blood vessel at a depth of 15 µm below the surface. The lines crossing the blood vessel walls in THG panels are plotted to the right with 0.4 µm per pixel sampling size. Scale bars in (a-b), 20 µm.
